# Reactive Oxygen Species Generation by Reverse Electron Transfer at Mitochondrial Complex I Under Simulated Early Reperfusion Conditions

**DOI:** 10.1101/2023.11.21.568136

**Authors:** Caio Tabata Fukushima, Ian-Shika Dancil, Hannah Clary, Nidhi Shah, Sergiy M. Nadtochiy, Paul S. Brookes

**Affiliations:** Departments of Anesthesiology, University of Rochester Medical Center; Departments of Biochemistry, University of Rochester Medical Center; Departments of Pharmacology and Physiology, University of Rochester Medical Center

**Keywords:** Mitochondria, Ischemia, Reperfusion, Complex-I, Reverse Electron Transfer, Reactive Oxygen Species

## Abstract

Ischemic tissues accumulate succinate, which is rapidly oxidized upon reperfusion, driving a burst of mitochondrial reactive oxygen species (ROS) generation that triggers cell death. In isolated mitochondria with succinate as the sole metabolic substrate under non-phosphorylating conditions, 90% of ROS generation is from reverse electron transfer (RET) at the Q site of respiratory complex I (Cx-I). Together, these observations suggest Cx-I RET is the source of pathologic ROS in reperfusion injury. However, numerous factors present in early reperfusion may impact Cx-I RET, including: (i) High [NADH]; (ii) High [lactate]; (iii) Mildly acidic pH; (iv) Defined ATP/ADP ratios; (v) Presence of the nucleosides adenosine and inosine; and (vi) Defined free [Ca^2+^]. Herein, experiments with mouse cardiac mitochondria revealed that under simulated early reperfusion conditions including these factors, overall mitochondrial ROS generation was only 56% of that seen with succinate alone, and only 52% of this ROS was assignable to Cx-I RET. The residual non-RET ROS could be partially assigned to complex III (Cx-III) with the remainder likely originating from other ROS sources upstream of the Cx-I Q site. Together, these data suggest the relative contribution of Cx-I RET ROS to reperfusion injury may be overestimated, and other ROS sources may contribute a significant fraction of ROS in early reperfusion.

## Introduction

The metabolite succinate is critical to the pathogenesis of ischemia and reperfusion (IR) injury, which plays a role in pathologies such as heart-attack and stroke ^1–3^. Succinate is widely observed to accumulate during tissue ischemia, both from aspartate and fumarate via the reversal of mitochondrial complex II (Cx-II), and from glutamate and α-ketoglutarate via canonical (clockwise) Krebs’ cycle activity ^1,2^. Regardless its mechanism, the highly conserved nature of succinate accumulation across diverse tissues and species ^4^ is consistent with beneficial effects during ischemia: Cx-II reversal permits continued operation of complex-I (Cx-I) proton pumping, while clockwise TCA cycle operation generates GTP via substrate-level phosphorylation, such that overall succinate accumulation can support ATP synthesis in the absence of oxygen ^2,5^. However, upon tissue reperfusion the rapid oxidation of succinate by mitochondrial Cx-II is a key driver of reactive oxygen species (ROS) generation ^1,6^, which along with Ca^2+^ overload triggers opening of the mitochondrial permeability transition (PT) pore and subsequent cell death ^7–10^.

Experiments incubating isolated mitochondria with combinations of metabolic substrates and inhibitors have led to the elucidation of at least 8 different sites for ROS generation within the organelle ^11^, with the most widely studied of these being the ubiquinone (Q) binding sites of respiratory complex I (Cx-I) and complex III (Cx-III). In mitochondria exposed to succinate as sole respiratory substrate and under classical state-4 (non-phosphorylating) respiration conditions, the bulk of ROS generation (>90%) can be blocked by the Cx-I inhibitor rotenone ^12–14^, thus revealing the phenomenon of reverse electron transfer (RET), in which electrons from the Q pool reduced by Cx-II can drive ROS generation at the Q binding site of Cx-I. Complex I RET ROS depends strongly on the trans-membrane potential ^14^, and a series of specific inhibitors of this phenomenon (S1QELs) have been developed as potential therapeutics for IR injury ^15,16^.

The combined observations of a key role for succinate oxidation in IR injury, and Cx-I RET as the main source of ROS when succinate is substrate, have led to the general assumption that Cx-I RET is *the* source of ROS during reperfusion ^1,2^. However, despite robust protection from IR injury by the S1QEL compounds ^16^, a focus on Cx-I RET as the only ROS source in IR may have overlooked other conditions in early reperfusion that could impact mitochondrial ROS generation. Herein, 6 such factors were examined.

During ischemia, mitochondrial respiratory inhibition leads to accumulation of NADH, a phenomenon that can be observed via NAD(P)H autofluorescence in ischemic cells and tissues ^6,17,18^. It is reasonable to assume that, at the time of reperfusion, the pyridine nucleotide pool is maximally reduced, and this would be expected drive Cx-I in its forward direction.

The metabolite lactate is well-known to accumulate in ischemic tissues as a result of anaerobic glycolysis, and could be a potential substrate for mitochondrial respiration ^19,20^. Recently, lactate has also been shown to impact mitochondrial function independent of its own metabolism ^21^. In addition to lactate itself (*pK*_a_ 3.9), glycolysis generates protons from ATP hydrolysis at hexokinase and phosphofructokinase, and along with protons from CO_2_ generated by the TCA cycle ^22–24^ this results in intracellular acidification to approximately pH 6.8 in the case of cardiac ischemia ^25,26^. We previously reported that acidic pH may stimulate Cx-I RET ROS, but simultaneously inhibits Cx-II activity, such that it may have no overall effect on succinate-driven Cx-I RET ROS ^27^. Nevertheless, how pH interacts with the other factors studied herein is unknown.

Another factor that may impact Cx-I RET ROS is the purine nucleotide pool. Although RET has mostly been studied in non-phosphorylating conditions (i.e., classical state-2 or state-4 respiration), it is virtually absent in actively phosphorylating mitochondria (classical state-3 in the presence of ADP) ^11,14^. The situation in early reperfusion is more nuanced. During ischemia the cessation of mitochondrial ATP synthesis leads to a precipitous drop in [ATP], while purine nucleotide breakdown also leads to lower [ADP], yielding AMP and the nucleosides adenosine and inosine ^28,29^. At the start of reperfusion, there is undoubtedly a non-zero amount of ADP present, such that mitochondrial respiration may be in an intermediate respiratory “state 3.5”, with some of the transmembrane potential being consumed for rapid ATP synthesis. In addition to the documented effects of lowered transmembrane potential on Cx-I RET ROS ^14^, nucleosides may also impact mitochondrial function via their effects on K_ATP_ channels ^30,31^.

Finally, due to the loss of ion homeostasis (see ^32,33^ for review), accumulation of both cytosolic and mitochondrial Ca^2+^ is a hallmark of ischemia. Although ultimately an overload of Ca^2+^ is thought to work in concert with ROS to trigger the mitochondrial PT pore ^7,8^, the impact of moderate amounts of Ca^2+^ during early reperfusion on Cx-I RET ROS is unclear.

Herein, using isolated heart mitochondria, we sought to build a model of simulated early reperfusion incorporating each the above factors, to determine their impact on Cx-I RET ROS. Our results suggest the contribution of Cx-I RET to ROS generation in reperfusion may be somewhat overestimated, but it nevertheless remains the major ROS source under such conditions and therefore remains a viable target for therapeutic intervention.

## Methods

C57BL/6J mice of both sexes were bred in-house and maintained on a 12-hr. light/dark cycle with food and water *ad libitum*. All experiments conformed with the US Guide for the Care and Use of Laboratory Animals and were approved by a local animal ethics committee (protocol #UCAR2007-087).

Mitochondria were isolated by protease digestion to sample both subsarcolemmal and interfibrillar populations ^34^. Following tribromoethanol anesthesia, the heart was chopped in ice-cold isolation media (“IM”) comprising (in mM): KCl (100), Tris (50), EGTA (2), pH 7.3. Tissue was suspended for 3 min. in 15 ml of IM supplemented with BSA (0.5% w/v), MgCl_2_ (5 mM), ATP (1 mM) and proteinase (2.1 U/mL, Sigma P8038), followed by homogenization (IKA Tissumizer, 21,000 rpm, 10 s.), a further 3 min. of mixing, dilution to 50 ml in IM, then centrifugation (600 x *g*, 5 min, 4°C). Supernatants were centrifuged (10,000 x *g*, 10 min, 4°C), and pellets subjected to 2 more wash steps (10,000 x *g*, 10 min, 4°C) before final suspension in 150 µL IM and storage in the dark on ice. Protein was determined using the Folin-Phenol reagent ^35^.

Mitochondrial H_2_O_2_ generation was measured using a Agilent/Varian Cary spectrofluorometer. The thermostated (37°C) cuvet holder was raised to facilitate a sample volume of 0.6ml, with the sample stirred by a motor-driven paddle. Mitochondria (0.5mg protein/ml) were incubated in buffer comprising: KCl (120), sucrose (25), MgCl_2_ (5), KH_2_PO_4_ (5), EGTA (1), HEPES (10), horseradish peroxidase (HRP, 2U/ml), superoxide dismutase (SOD, 80 U/ml), *p*-hydroxyphenylacetic acid (*p*HPA, 100 µM), fat-free bovine serum albumin (0.1 % w/v), pH 7.4 or 6.8 as required. For PCr/Cr clamp experiments, HEPES was at 30 mM to more strongly buffer pH, and KCl was at 80 mM to maintain osmolarity due to added Na^+^ (see below). Aliquots of buffer solutions were stored at -20°C, and each aliquot re-frozen no more than once. Excitation/emission wavelengths used for *p*HPA were 320/410 nm respectively, with a slit-width of 5 nm. Readings were taken for 0.5 s. every 2 s. (25% duty cycle), with the photomultiplier set at 600 volts.

PCr/Cr ratios were selected assuming *K_eq_* = 150 for the reaction: *ADP + PCr ↔ ATP + Cr* ^36,37^. Reported ATP/ADP ratios in the heart are subject to considerable variation, but generally fall in the range 100-300 ^38–41^, with more recent non-invasive methods such as *in-situ* ^31^P-NMR yielding higher ratios than “grind-and-find” methods such as HPLC wherein the labile nature of these metabolites may be impacted by extraction ^42^. Although fluorescent ATP/ADP biosensors are available ^43^, even newer variants achieve maximal signal at ATP/ADP ratios of 10-fold ^44^, far below those seen in the heart. Work from ours and other labs suggests that ATP/ADP drops by up to 4-fold during ischemia ^26,39,45^. As such, we chose three different ATP/ADP ratios to study: 3000/1, representing excess ATP and mimicking state-4 respiration, 145/1 as an intermediate state approximating the normal heart, and 60/1 representing an ischemia-like condition, cognoscent of the rapid recovery of ATP synthesis upon tissue reperfusion.

Creatine monohydrate, phosphocreatine disodium, and NaCl were added as per the table below, maintaining total creatine species at 20mM and total Na^+^ at 40 mM. Although recent work has implied a role for sodium in the regulation of mitochondrial ROS generation from Cx-III ^46^, herein no significant effect of 50 mM NaCl on Cx-I RET ROS was observed (data not shown). ADP was initially added at 150 µM, and creatine kinase (CK) at 15 U/ml. As such, our ischemia-like condition would result in a nominal [ADP] of 2.5 µM.

| *Condition* | *ATP/ADP* | *PCr/Cr* | *PCr (mM)* | *Cr (mM)* | *NaCl (mM)* |
| --- | --- | --- | --- | --- | --- |
| State-4-like | 3000/1 | 19/1 | 19 | 1 | 2 |
| Intermediate | 145/1 | 1/1 | 10 | 10 | 20 |
| Ischemic | 60/1 | 2/5 | 5.7 | 14.3 | 28.6 |

As necessary during measurements, additions to the assay included: succinate (4 mM), pyruvate (5 mM), carnitine (5 mM), lactate (30 mM), adenosine (1.7 mM) or inosine (2 mM). 30mM lactate was chosen as being in the pathologic range ^47^ while also being higher than the level of pyruvate used, to maintain a lactate/pyruvate ratio appropriate for the heart^48^. Concentrations of nucleosides were chosen based on several prior observations in ischemic heart tissue ^47,49–53^, with appropriate corrections applied for conversion between units ^54^. All additions were from 100x stocks in the same pH-corrected buffer. Inhibitors added included rotenone (3.3 µM), S1QEL-1.1(1 µM) ^15,16^, or S3QEL-2 (3.3 µM) ^55^, from 100x stocks in DMSO. The SnQEL compounds were sonicated immediately prior to use, due to solubility issues experienced with frozen stocks. At the end of each run, traces were calibrated by 2 additions of 1 nmol freshly prepared H_2_O_2_.

For Ca^2+^ experiments, free [Ca^2+^] was buffered by EGTA (∼1 mM), and concentrations calculated using a version of the MaxChelator web-app ^56^ that takes into account prevailing pH, [ATP] and [Mg^2+^]. Free Ca^2+^ concentrations of 5 and 12 µM were chosen as representing a clear elevation from baseline levels in the heart (0-1 µM), while avoiding the high concentrations known to acutely trigger PT pore opening (20-100 µM) ^57^.

In figures, N refers to biological replicates, with each N being a separate measurement made on an independent mitochondrial preparation from a single animal. Statistical significance for differences between experimental groups was determined via ANOVA followed by two-tailed Student’s *t*-test, with α set at 0.05. Due to the complex interplay between the factors studied here, each could not be studied independently or in all 36 possible combinations. Rather, each stage of the experimental model built on the previous stage, adding a new factor each time in the order: NADH, lactate, pH 6.8, ischemic ATP/ADP, purines, Ca^2+^.

## Results

The complete original data set for this paper is archived online at FigShare (DOI: 10.6084/m9.figshare.24585234).

### Baseline condition – Succinate alone drives Cx-I RET ROS

The data in Figure 1 confirm previous reports ^12–14^, showing mitochondrial ROS generation in the presence of succinate alone is largely inhibited by the Cx-I inhibitor rotenone (**Figure 1A/B**). In addition, succinate-driven ROS was inhibited by the specific Cx-I Q site ROS inhibitor S1QEL, with no further impact seen upon addition of the Cx-III Q site ROS inhibitor S3QEL (**Figure 1C/D**). Thus, under these conditions of succinate alone and non-phosphorylating respiration, 89% of ROS generation is attributable to Cx-I RET. Note that small deflections in the fluorescent signal observed upon addition of S1QEL and S3QEL were likely caused by light scatter due to poor solubility of these compounds (see methods).

**Figure 1.**
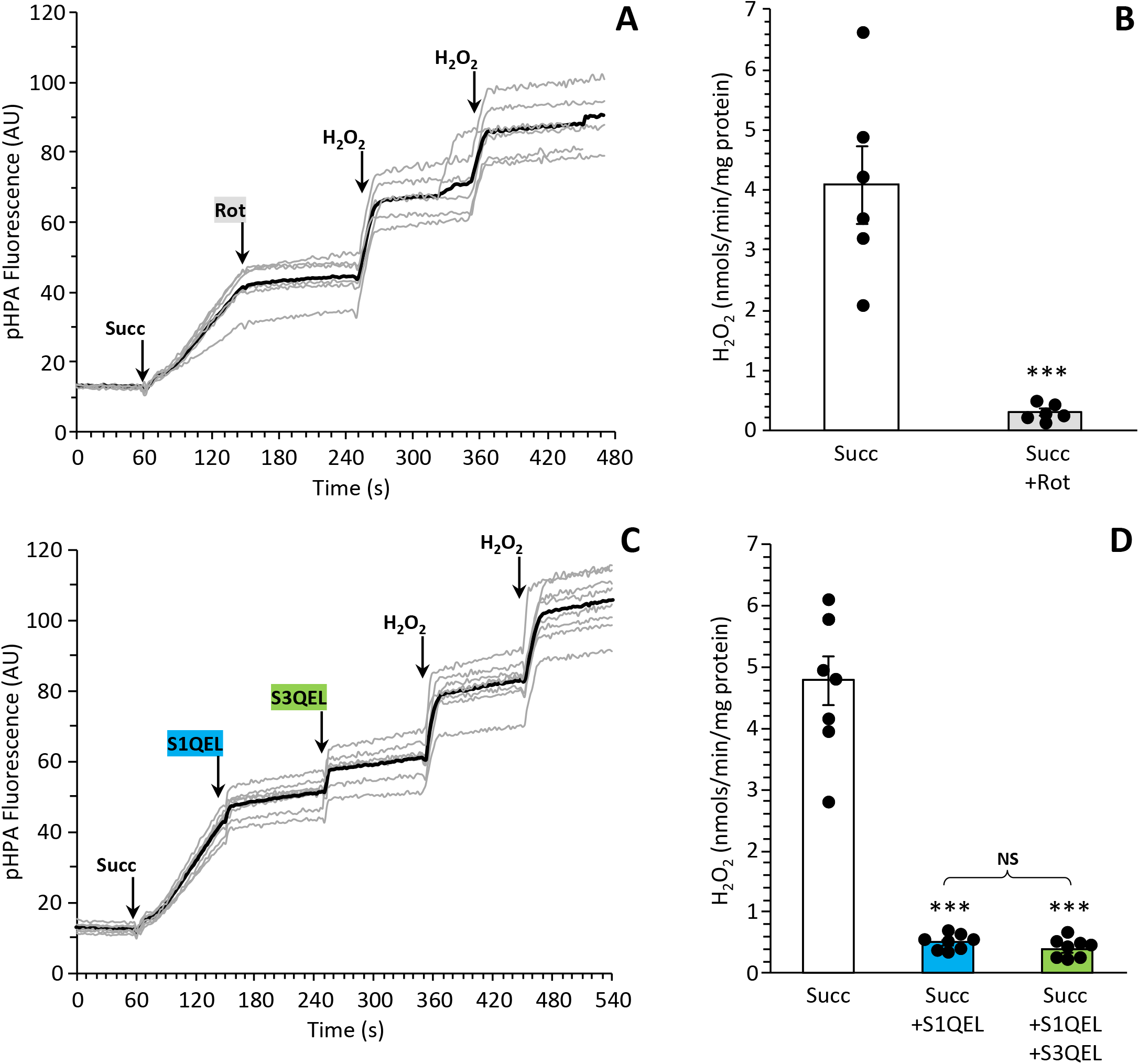
Baseline ROS Generation by Cx-I RET. Mouse heart mitochondria were incubated as described in the methods, with ROS generation measured by pHPA fluorescence. **(A):** *p*HPA fluorescent traces (individual traces in gray, average in black). Where indicated, succinate, rotenone, and H_2_O_2_ were added. **(B):** Quantitation of H_2_O_2_ generation rates from traces as shown in A. Individual data points are superimposed on bars showing means ± SEM. **(C):** As in A, but with use of S1QEL and subsequently S3QEL instead of rotenone. **(D):** As in B. ***p<0.0005 vs. baseline condition. NS: not significant.

### Effect of NADH

At the start of reperfusion both [NADH] and [succinate] are expected to be high. To avoid complications from the position of Cx-II within the TCA cycle, we sought to generate NADH in a manner that did not engage the cycle. To accomplish this, mitochondria were incubated with pyruvate plus carnitine, to generate NADH at pyruvate dehydrogenase (PDH), with the subsequent export of acetyl-carnitine serving both to relieve feedback-inhibition on PDH, as well as prevent acetyl-CoA transmission into the TCA cycle (**Figure 2A**) ^58,59^. Fluorescent measurement of NAD(P)H (**Figure 2B**) showed that pyruvate alone led to partial reduction of the pyridine nucleotide pool, with subsequent addition of carnitine fully reducing it (due to relief of PDH inhibition). Addition of ADP oxidized the pool, indicating appropriate coupling of oxidative phosphorylation, and inhibition of Cx-I by rotenone again fully reduced the pool.

**Figure 2:**
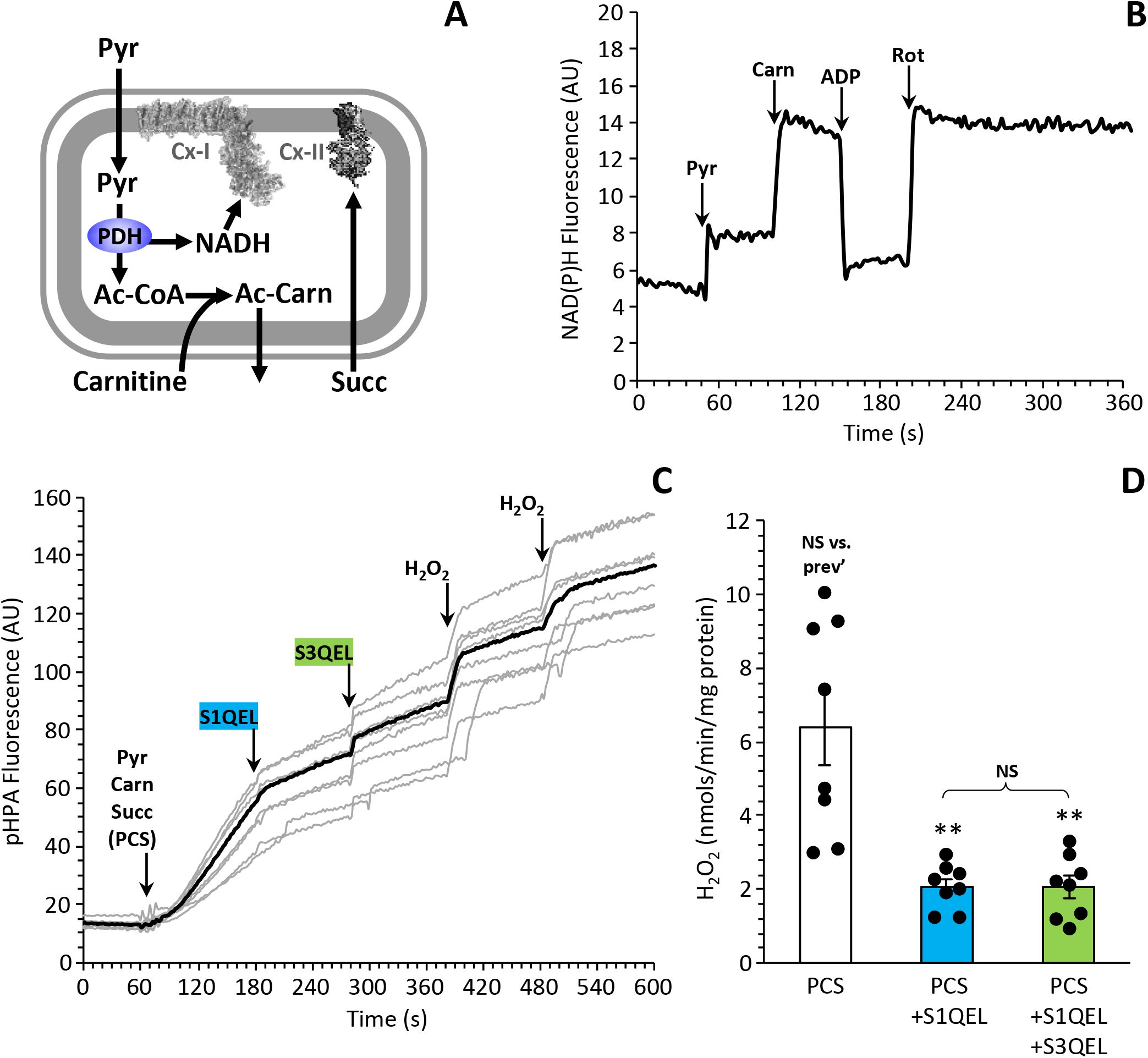
Impact of NADH on ROS. **(A):** Schematic showing the incubation system to generate NADH from PDH, with export of acetyl-CoA from mitochondria by carnitine, thus avoiding engagement of the TCA cycle. **(B):** NADH fluorescence in isolated mitochondria (λ_EX_ 340nm /λ_EM_ 460nm), with sequential additions of pyruvate, carnitine, ADP, and rotenone. Representative trace from 3 similar experiments. **(C):** *p*HPA fluorescent traces (individual traces in gray, average in black). Where indicated, pyruvate, carnitine, succinate, S1QEL, S3QEL and H_2_O_2_ were added sequentially. **(D):** Quantitation of H_2_O_2_ generation rates from traces as shown in C. Individual data points are superimposed on bars showing means ± SEM. **p<0.005 vs. baselne condition. NS: not significant.

As shown in **Figure 2C/D**, the combination of pyruvate, carnitine and succinate (PCS) drove ROS generation at a rate 33% higher than succinate alone (compare to **Figure 1D**). A direct comparison between all of the conditions studied is shown in **Figure 6A**, indicating that this additional ROS was not due to Cx-I RET or Cx-III, since it was not impacted by the inhibitors S1QEL or S3QEL. As such, it is likely this additional ROS originates either from PDH or from the upstream flavin site in Cx-I ^60^.

### Effect of lactate and acidic pH

Lactate is greatly elevated in ischemic tissues, and its potential use as a fuel source by a variety of tissues has been the subject of intense interest ^19^. Herein, **Figure 3A/B** shows that addition of 30 mM lactate onto the previous experimental condition (**Figure 2C/D**) did not alter the pattern or magnitude of ROS generation.

**Figure 3:**
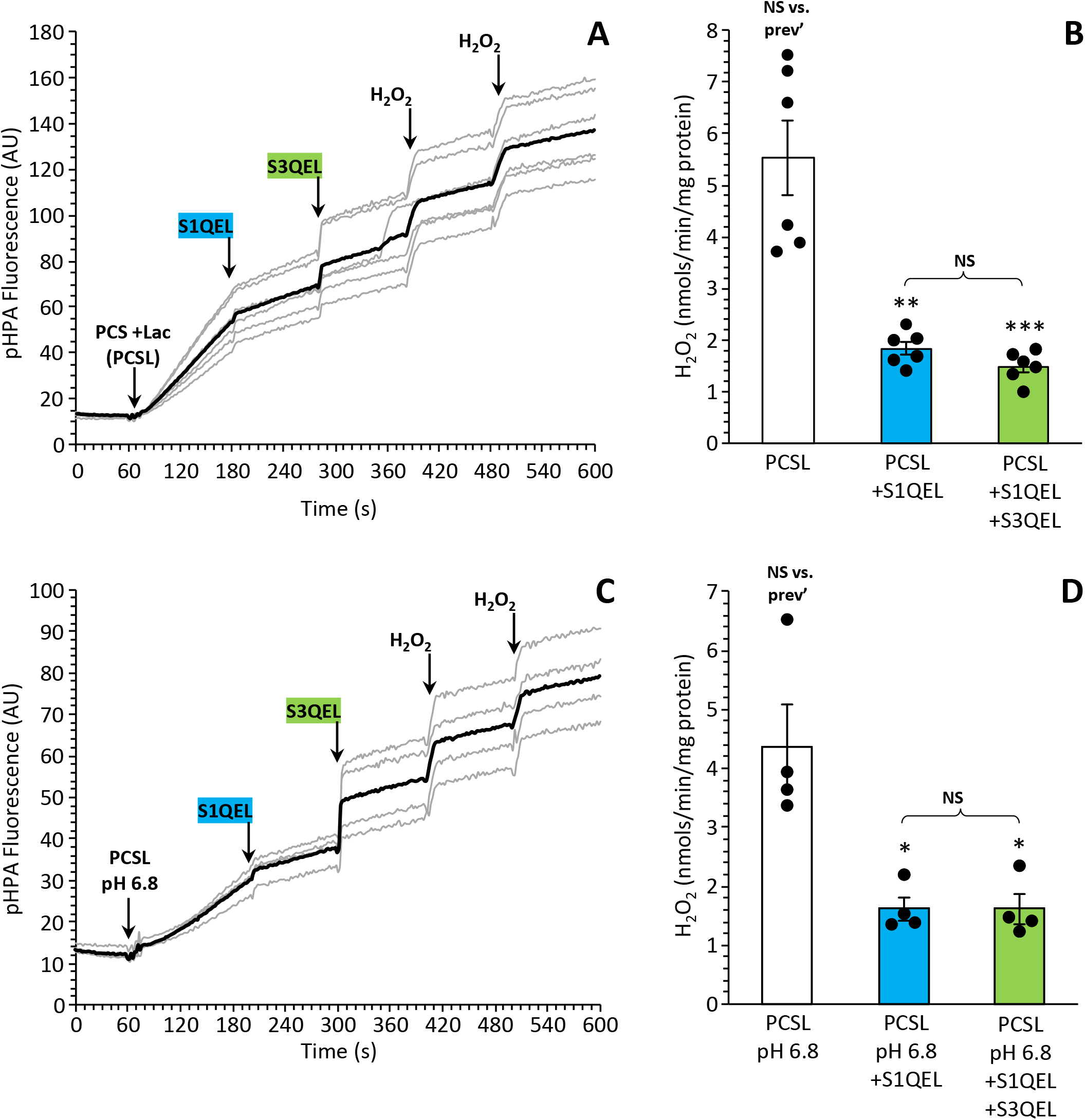
Impact of Lactate and Acidic pH on ROS. **(A):** *p*HPA fluorescent traces (individual traces in gray, average in black). Where indicated, pyruvate, carnitine, succinate, lactate, S1QEL, S3QEL and H_2_O_2_ were added sequentially. **(B):** Quantitation of H_2_O_2_ generation rates from traces as shown in A. Individual data points are superimposed on bars showing means ± SEM. **(C):** As in A, but at pH 6.8. **(D):** As in B. *p<0.05, **p<0.005, vs. baseline condition. NS: not significant.

Considering the impact of acidic pH, **Figure 3C/D** shows that imposition of pH 6.8 onto the lactate condition did cause a 21% drop in overall ROS generation (**see** **Figure 6A**), with the bulk of this decrease being in Cx-I RET (down 25%), while the contribution of other sources remained largely intact. This finding is consistent with our previous results, showing that even though acidic pH may stimulate RET itself, overall it inhibits succinate-driven RET ROS via inhibition of Cx-II activity ^27^. Importantly (and as previously reported ^27^), no impact of pH was seen on the ROS measurement system, and it should be noted that even if such an effect existed it would be corrected by the use of internal calibration with authentic H_2_O_2_ in every trace.

### Effect of ATP/ADP ratio and purines

The dependence of Cx-I RET ROS on the mitochondrial transmembrane potential ^14^ means that it is typically studied under non-phosphorylating (state-4) conditions, with addition of bolus ADP to initiate state-3 respiration effectively eliminating the phenomenon. However, the impact of physiologically relevant ATP/ADP ratios on Cx-I RET ROS is unclear. Herein, we employed a PCr/Cr/CK clamp to buffer ATP/ADP ratios (**Figure 4A**). Three different ATP/ADP conditions were selected: state-4 like (3000/1), ischemia-like (60/1), and intermediate (145/1). This system was built onto the previous stage of the model (i.e., pyruvate, carnitine, succinate, lactate, pH 6.8, **Figure 3C/D**) as a baseline condition.

**Figure 4:**
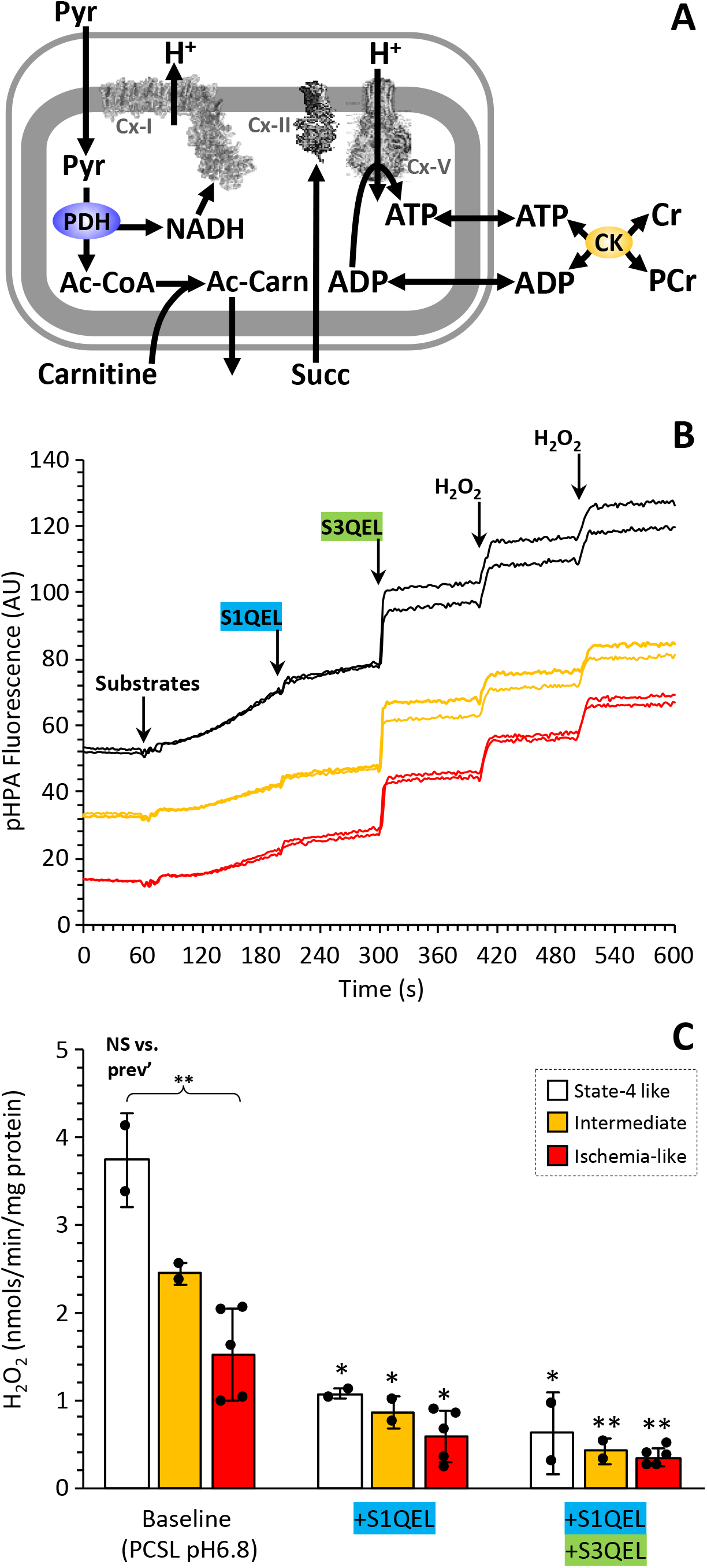
Impact of Ischemia-Like ATP/ADP on ROS. **(A):** Schematic showing the creatine kinase (CK) and phosphocreatine/creatine (PCr/Cr) buffer system. **(B):** *p*HPA fluorescent traces. Where indicated, substrates, S1QEL, S3QEL and H_2_O_2_ were added sequentially. Two independent traces are shown for each ATP/ADP condition (color coded), with the data offset vertically by 20 units for clarity. **(C):** Quantitation of H_2_O_2_ generation rates from traces as shown in B. Individual data points are superimposed on bars (color-coded to match B) showing means ± SEM. Individual data points are superimposed on bars showing means ± SEM. *p<0.05, **p<0.005, vs. baseline condition. NS: not significant.

As shown in **Figure 4B/C**, the pattern of ROS generation under state-4 like conditions largely echoed that seen in true state-4 (no ADP) in **Figure 3C/D**. However, decreases in the ATP/ADP ratio had a profound impact on ROS generation, with the ischemia-like condition resulting in a 65% drop in overall ROS (see comparison **Figure 6A**). Notably the contribution of Cx-I RET ROS remained fairly constant at 62% of the total, suggesting that all sources of ROS were impacted by the imposition of a lower ATP/ADP ratio. Taking this condition as a new baseline, **Figure 5A** shows that further addition of the nucleosides adenosine and inosine did not impact the overall pattern or magnitude of ROS generation.

**Figure 5:**
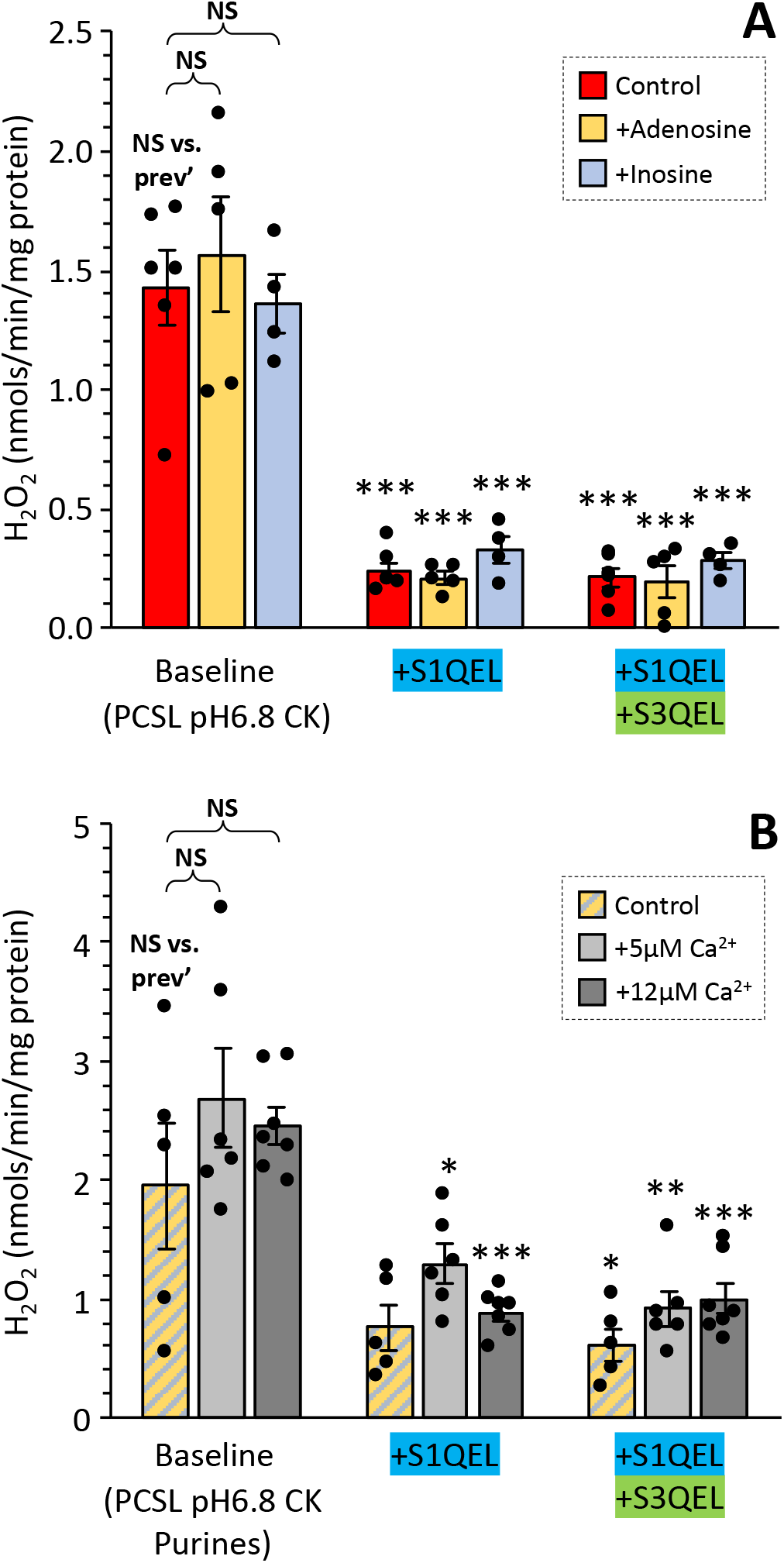
Impact of Purine Nucleosides and Ca^2+^ on ROS. **(A):** Effect of adenosine or inosine on ROS generation (original *p*HPA traces not shown). Graph shows rates of H_2_O_2_ generation calculated from traces, with the ischemia-like ATP/ADP condition from previous figure (red bar) forming the baseline condition for addition of nucleosides. **(B):** Quantitation of H_2_O_2_ generation rates under different prevailing free [Ca^2+^] conditions. The baseline condition herein represents the presence of both adenosine and inosine from panel A. Individual data points are superimposed on bars showing means ± SEM. *p<0.05, **p<0.005, ***p<0.0005, vs. baseline condition. NS: not significant.

### Effect of Ca^2+^

Both ischemia and reperfusion are characterized by derangements in cellular ion handling ^32^, and while the role of excess Ca^2+^ as a trigger for mitochondrial PT pore opening in reperfusion has been extensively studied ^7,8^, the impact of moderate Ca^2+^ levels experienced during ischemia on mitochondrial ROS is less well understood. Herein, we used an EGTA buffer system to impose two intermediate Ca^2+^ levels on the baseline condition from the previous stage of the model (i.e., with purines, **Figure 5A**). As shown in **Figure 5B**, both 5 µM and 12 µM free Ca^2+^ caused an elevation in overall ROS generation, which was not attributable to a stimulation of either Cx-I RET ROS or Cx-III ROS (i.e., the extra ROS was not sensitive to S1QEL or S3QEL – see **Figure 6A** for comparison).

**Figure 6.**
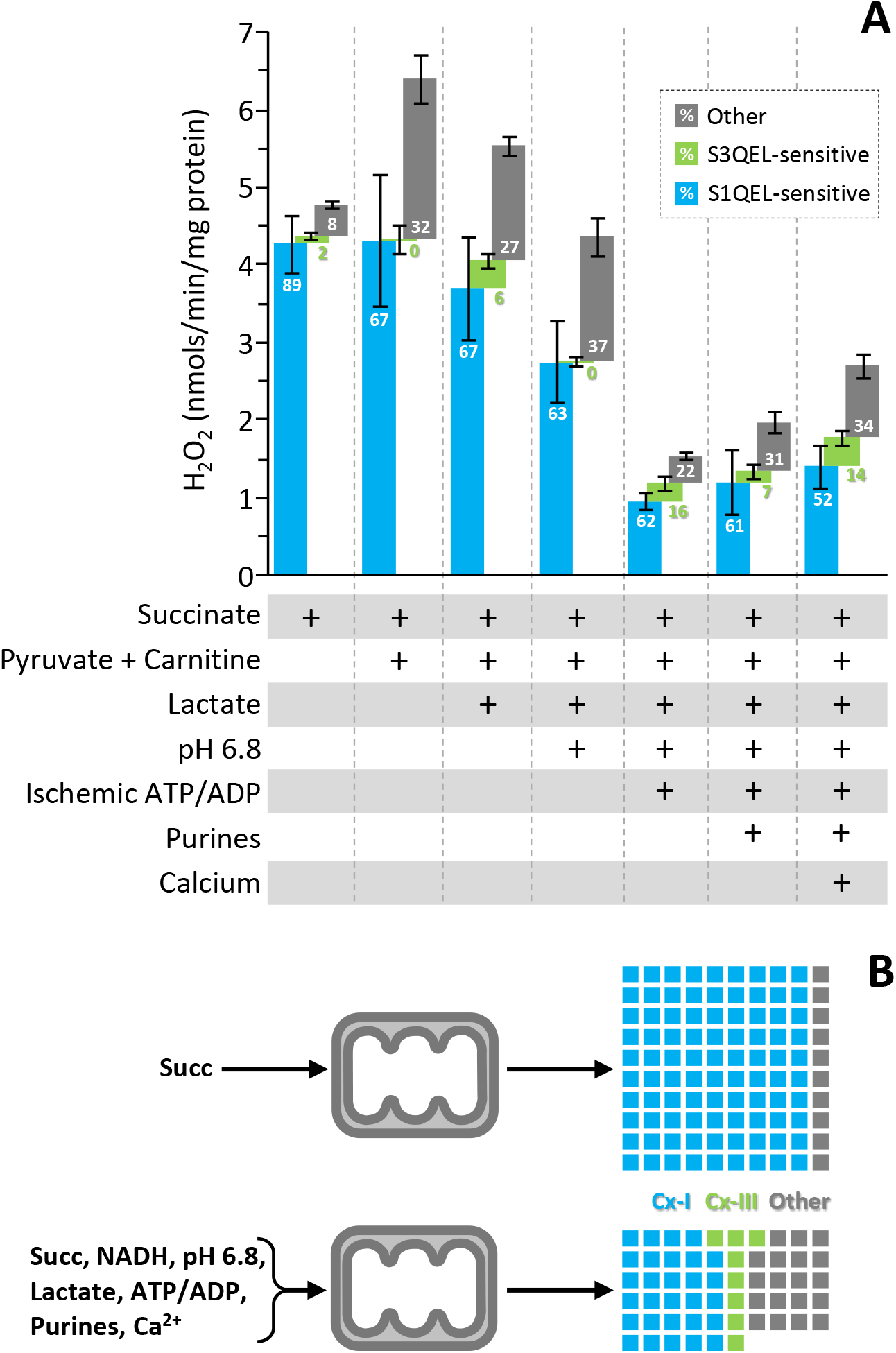
Overall Comparison of Conditions. **(A):** Comparison of all conditions studied herein, with developments to the model system proceeding left-to-right. Blue bars show S1QEL-sensitive component of ROS, green the S3QEL-sensitive, and gray the component not attributable to either source. The bars show absolute values of H_2_O_2_ (see Y-axis) with standard errors, while numbers within the bars show the percentage contribution of each component to the total. **(B):** Visualization of the overall and contributing components of ROS generation in the original (succinate only) and final (simulated early reperfusion) conditions. Each colored box represents 1 unit of ROS, with the original condition assigned to 100 units. In the final model, overall ROS is 56% of that seen in the original model. Contribution of each component is color-coded as per panel A, and % contributions scaled appropriately.

## Discussion

Comparing all the conditions studied herein (**Figure 6A**), the pattern of ROS generation seen under ischemia-like conditions (i.e., pyruvate, carnitine, succinate, lactate, pH 6.8, ischemia-like ATP/ADP, adenosine, inosine, Ca^2+^) is somewhat different from the succinate-only non-phosphorylating condition upon which much of the literature on IR injury is based. Specifically comparing the original and final models, as illustrated in **Figure 6B**, the overall ROS generation is 45% lower, and the contribution of Cx-I RET ROS to this total falls from 89% to 52%. Furthermore, the contribution of Cx-III ROS increases from essentially zero to 14% of the total, while additional unknown sources of ROS rise from 8 to 34% of the total. As such, it is concluded that the contribution of Cx-I RET ROS to ischemia-reperfusion injury may be over-estimated, and that of other ROS sources including Cx-III may be under-estimated.

The presence of a high concentration of NADH in addition to succinate appears to invoke ROS generation by an additional source (**Figure 2C/D**). Although it has recently been shown that S1QEL is capable of inhibiting ROS generation at the Cx-I Q site regardless the direction of electron flow through the complex ^61^, the additional ROS seen here with NADH was not S1QEL or S3QEL sensitive, ruling out the Cx-I or Cx-III Q sites as its source. As such, we speculate this additional ROS may originate from a site downstream of pyruvate, such as pyruvate dehydrogenase or the Cx-I flavin site ^60^.

The lack of effect of lactate on the overall pattern of ROS generation (**Figure 3A/B**) is likely due to the redox equilibrium of lactate dehydrogenase (lactate + NAD^+^ ↔ pyruvate + NADH + H^+^). There is debate regarding the presence of an isoform of LDH within mitochondria ^19,48^, but if present this enzyme would be prevented from converting lactate to pyruvate by the high prevailing [NADH] and [H^+^] (acidic pH). Accordingly, experiments performed with lactate as the sole substrate resulted in no ROS at all (data not shown).

While the apparent lack of effect of the nucleosides adenosine and inosine (**Figure 4D**) cannot rule out an effect of these metabolites in isolation, any effect of these metabolites may be masked by the large number of perturbations already present in the experimental system. In this respect, it is important to acknowledge a key limitation of this study: the use of a progressive experimental system wherein each experiment established a new baseline condition from which to add the next perturbation. Although the overall number of perturbations studied was small (**Figure 6A**), studying each of these independently and in all possible 36 combinations would be excessive (note that the buffer system for the Ca^2+^ experiments in **Figure 5B** alone comprised 20 components - KCl, sucrose, MgCl_2_, KH_2_PO_4_, EGTA, HEPES, HRP, SOD, *p*HPA, succinate, pyruvate, carnitine, lactate, CK, PCr, Cr, NaCl, adenosine, inosine, CaCl_2_). An additional limitation was our use of steady-state conditions, whereas tissue reperfusion is a dynamic system that changes rapidly. Indeed, measurements in intact hearts show elevated ROS generation within the first 5 seconds of reperfusion ^6^, while most succinate is consumed within 2 min., and NADH returns to baseline levels within 30 s. As such, accurate values for many parameters during early reperfusion are not available, so we based our conditions on values in late ischemia, being as close as possible to the start of reperfusion.

Several experimental phenomena discovered during these experiments but not shown in the results warrant brief discussion. Firstly, the SnQEL drugs caused an immediate increase in fluorescent signal, even in buffer alone. This was attributed to light scatter due to poor solubility and could be somewhat alleviated by sonication of stocks prior to use (see methods). Secondly, our choice to use *p*HPA instead of a more *modern* probe such as Amplex red ultra as we have used previously ^27^ was driven by an observation that carnitine interfered with the fluorescent signal of the latter (not shown). Thirdly, in the ATP/ADP experiments (**Figure 4B**), a significant lag was seen in the ROS slope following substrate addition. We attribute this to the fact that only ADP was present at the start of the experiment, and the PCr/Cr/CK system takes some time to reach equilibrium *in-situ*, bringing [ADP] down to a level that permits reestablishment of the transmembrane potential and ROS generation from Cx-I RET. Lastly, although it has been reported that inorganic phosphate (Pi) inhibits RET ^14^, we saw no such inhibition here with 5 mM Pi present in all buffers, and others have observed RET in the presence of 10 mM Pi ^12^.

Overall, our findings (**Figure 6B**) suggest that Cx-I RET ROS may be less important for the pathology of IR injury than previously thought, while the Cx-III Q site ^62^ and other ROS generating sites upstream of the Cx-I Q site (e.g., PDH and/or Cx-I flavin) may be more important than currently acknowledged. There are, of course, additional factors that could contribute to the mechanisms of ROS generation in the respiratory chain, including metabolites that accumulate in hypoxia such as L-2-HG, and defects in other metabolic pathways such as glycolysis or the pentose phosphate pathway ^26,63^. Nevertheless, in its current state our model of early-reperfusion-like conditions represents a baseline for future studies on this important pathologic phenomenon. Importantly, although the contribution of Cx-III ROS appears to be more than previously described (∼15% of total ROS), the bulk of ROS (over 50%) still originates from Cx-I RET, highlighting the importance of this site as a therapeutic target. Furthermore, the importance of succinate as a substrate for driving ROS at reperfusion is not impacted by these studies (both Cx-I RET and Cx-III are downstream of succinate), so therapeutic efforts to limit succinate oxidation at reperfusion are still very much valid.

## Acknowledgements

This work was funded by a grant from the US National Institutes of Health (R01-HL071158) to PSB. We thank Alex Milliken, PhD (URMC) for technical assistance, and for critical reading of the manuscript.

